# Identification of Anonymous DNA Using Genealogical Triangulation

**DOI:** 10.1101/531269

**Authors:** Paul Ellenbogen, Arvind Narayanan

## Abstract

Consumer genetics databases hold dense genotypes of millions of people, and the number is growing quickly [1] [2]. In 2018, law enforcement agencies began using such databases to identify anonymous DNA via long-range familial searches. We show that this technique is far more powerful if combined with a genealogical database of the type collected by online ancestry services. We present a “genealogical triangulation” algorithm and study its effectiveness on simulated datasets. We show that for over 50% of targets, their anonymous DNA can be identified (matched to the correct individual or same-sex sibling) when the genetic database includes just 1% of the population. We also show the effectiveness of “snowball identification” in which a successful identification adds to the genetic genealogical database, increasing the identification accuracy for future instances. We discuss our technique’s potential to enhance law enforcement capabilities as well as its privacy risks.

## Introduction

In 2018, US law enforcement agencies began using long-range familial searches to identify perpetrators based on crime-scene DNA. This works as follows: investigators obtain a dense genome-wide genotyping profile from the DNA, masquerade it as a profile obtained from a consumer genetics provider (such as 23andme or AncestryDNA), and upload it to a genetic genealogy service to find genetic relatives. In a typical successful case, a single match of a 2nd or 3rd cousin might be found, resulting in a few tens or hundreds of possibilities for the perpetrator’s identity. Investigators then use age, sex, geography, and other clues to prune the list. Recently, Erlich et al. estimated the effectiveness of this technique using genomic data of 600,000 individuals, concluding that half the searches of European-descent individuals will result in a third-cousin-or-closer match [3]. They also estimate the efficacy of pruning.

A major limitation of the current approach is the need for time-consuming manual pruning of a set of possibly hundreds of candidates, potentially requiring days or even months of police work [4][5][6]. Our technique builds on the existing approach, but automates the entire identification process and eliminates the need for manual pruning. We achieve this in a setting where a genealogical database is also available to the analyst, such as via an online ancestry service. We consider the case where a search of the genetic database yields multiple relatives (as defined by the presence of Identical By Descent (IBD) segments), however distant. We believe that this is already the case today on commercial genetic genealogy services such as 23andme for the majority of European-descent individuals. Each match yields a set of possibilities for the identity of the unknown individual; this allows the analyst to narrow the identity of the unknown individual to the intersection of these sets (Figure 1). This “genealogical triangulation” technique has the potential to yield a unique sibship.

**Figure 1:**
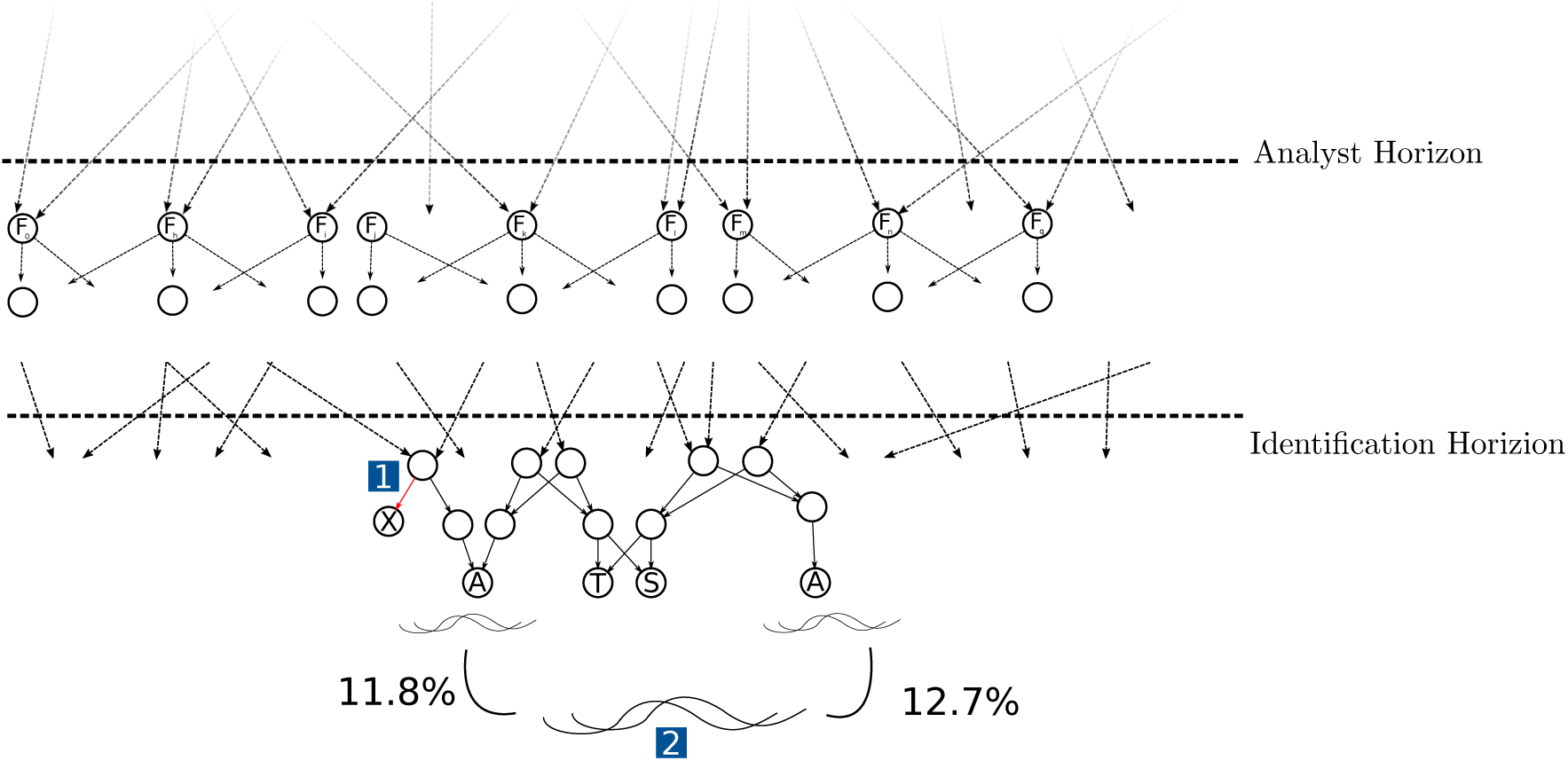
Overview of our setting and approach. The few most recent generations of the genealogy of the population are available to the analyst, albeit with missing or erroneous links (1). The analyst has dense genotype data for a small fraction of nodes in the tree, called anchors (denoted A). The analyst aims to find the node associated with an anonymous piece of DNA (2); the node is known to be below the identification horizon. The analyst begins by computing the IBD between the target genome and each anchor genome. In this example, the sibling group {*S, T*} matches the observed IBD of approximately 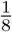 with both anchor nodes. *X* also matches the observed IBD for one anchor node, but not for the other. The analyst concludes that the DNA belongs to either *S* or *T*. Unlike this toy example, the closest IBD matches in a realistic setting are much more distant (e.g. fourth or fifth cousins). To estimate the probability distribution of IBD between any two nodes in the graph, the analyst makes the simplifying assumption that the founders (*F*_*1*_,*…, F*_*n*_) are unrelated, and simulates meiosis.

**Figure 2:**
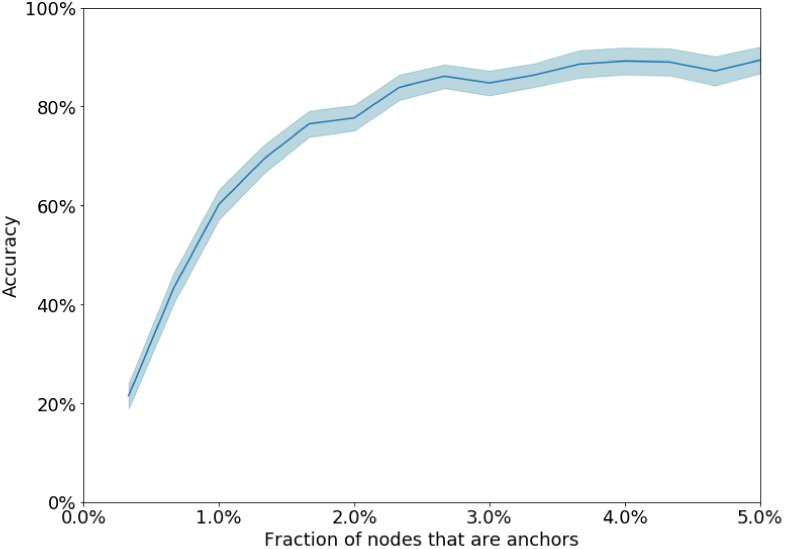
Identification accuracy for standard parameters.

We find that the availability of dense genotype data for as little as 1% of the population results in accurate identification in the median case, even in the presence of some error and incompleteness in the genealogy. Our technique is especially effective for en-masse reidentification of a large database of de-identified genotypes, as successful identification of a record allows the analyst to augment the genetic genealogical database, increasing the identification accuracy for the remaining records. This en-masse reidentification, which we call snowball identification, is made possible by the fact that our algorithm eliminates the manual pruning step.

Our results are obtained using synthetic data by simulating mating and recombination for populations of up to 100,000 individuals per generation. We use synthetic rather than actual data because comprehensive genomic data paired with genealogical data has not been released to the public as of this writing. Synthetic data are frequently used in population genetics studies [7]. While we do not model existing population structure, we simulate a variety of mating and migration models and show that our findings are broadly applicable.

## Methods

We developed two algorithms, simulation and identification, that will be deployed by the analyst. The simulation algorithm takes as input a set of population parameters, and simulates meiosis in such a population starting from a set of founders. The identification algorithm takes as input a set of information assumed to be available to the analyst, together with an anonymous genotype, and outputs the node that is the most likely match in the population.

There are two views of our synthetic universe: a god’s-eye view and an analyst view. The god’s eye view is the true state of our synthetic world, with all genealogical data and genomic data assumed to be correct and complete. The analyst’s view contains incomplete or erroneous genomic and genealogical information, and reflects the limitations of the data available to a real-world analyst. Thus, the set of genomes available to the analyst will be a (small) subset of the whole populations’ genomes. Additionally, IBD detection is an error prone process for the analyst, which we model via a threshold segment length beneath which the analyst cannot detect IBD segments. In the analyst’s view, the IBD between pairs of nodes in the graph is a probability distribution. Estimating this distribution is one of the main goals of simulation.

To generate the god’s eye view, we first generate genotypes for the population founders (note that the god’s eye view population extends above the analyst horizon), with the assumption that none of the founders are related, then propagated through the descendants. Starting from the founders, we generate genotypes of successive generations using a forward-in-time simulation technique. We track haplotypes in a similar way to the work of Kessner et al [8]. To simulate meiosis, we use recombination rates provided by the HapMap project [9].

The analyst’s simulation uses exactly the same technique, but on his view rather than the god’s eye view. The analyst repeats the simulation hundreds of times; each run produces a sample of the IBD for each pair of individuals. The analyst models IBD as a hurdle gamma distribution^1^ and estimates its parameters (for each pair of individuals) using the simulation outputs. The analyst is aware that this view of the genealogy is inaccurate and incomplete, and therefore expects the presence of cryptic IBD, that is, IBD segments between seemingly unrelated pairs of individuals. He estimates the distribution of cryptic IBD lengths by computing IBD among a sample of unrelated anchor nodes.

### Algorithm 1: Simulation and estimation algorithms (analyst view).

Given: a population genealogy and dense genotype data corresponding to a subset of nodes (anchors)

Fix a sufficiently large sample size *S*

Simulate genomes of the entire population *S* times using forward-in-time simulation

For each node *x* and each anchor node *a* ≠*x*:

Fit a hurdle gamma distribution *ξ*_*x,a*_ to the set of *S* simulated IBD lengths between *x* and *a*

Output the parameters of *ξ*_*x,a*_

Sample *S* pairs of anchor nodes 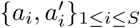 such that *a*_*i*_ and 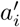 are unrelated in the genealogy

Fit a hurdle gamma distribution *ξ*_*cryptic*_ to the set of *S* observed IBD lengths between *a*_*i*_ and 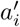

Our population model has a number of parameters, including monogamy rates, island configuration, and migration rates (1). Additionally, for the analyst’s view, the rates of missing data and incorrect data are configurable. We pick default values for these parameters based on empirical data about typical real-world values (to the extent that such data are available). To test the robustness of genealogical triangulation, we repeat our analysis with different values for each of the configurable parameters.

### Algorithm 2: Identification.

Given: a population genealogy, dense genotype data of anchors, and an anonymous dense genotype record.

Let

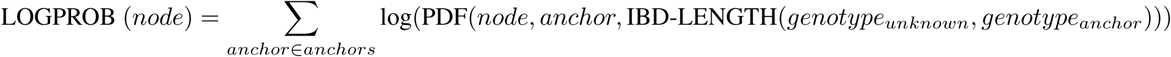

where

PDF(*x, y, ibd*_*length*)) = *ξ*_*x,y*_(*ibd*_*length*) if *x* and *y* are genealogically related and *ξ*_*cryptic*_(*ibd*_*length*) otherwise

Let *N* = 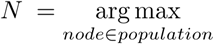 LOGPROB (*node*)

Output the sibship containing the node *N*.

Our identification algorithm is a Naive Bayes classifier which aims to predict which of a set of nodes in a genealogical graph is the best match for an anonymous genome. It starts with a uniform prior probability distribution over the entire population. The features are the IBD segments between the anonymous genome and each of a set of anchors: individuals in the genealogical graph whose genotype data are available to the analyst. For each anchor, the algorithm considers only the total IBD length represented by the segments (and not, for example, the number of segments).

For each population and error model interest, we evaluate our the accuracy of our algorithms as follows:

1. Synthesize the god’s eye view of the genealogy based on the population model
2. Synthesize the god’s eye view of the genotypes based on forward-in-time simulation
3. Synthesize the analyst view of the genealogy from the god’s eye view based on the error model
4. Copy the genomes of the anchors to the analyst view
5. Run the simulation and estimation algorithms for the analyst view
6. Run the identification algorithm on genotypes randomly sampled non-anchor nodes and evaluate the accuracy.

For each population and error model, there is a trade-off between the number of anchors and the probability of correct identification of a genome belonging to a random individual from the population. Our experiments seek to quantify this tradeoff. We consider identification successful if the identified sibship contains the true individual. A variant of our identification algorithm produces two outputs that might further aid the analyst. First, it produces a confidence score (see Supplement), which can be used to tune the sensitivity and specificity of identification. Second, it produces a ranked list of candidates based on the log-probability scores in Algorithm 2. Recall that an analyst might use further clues such as age and geography to prune such a list, which are beyond the scope of our research. In our experiments, we quantify the sensitivity-specificity trade-off as well as the usefulness of ranked shortlists of various sizes.

Finally, we introduce “snowball identification”, wherein we assume that the analyst has access to a small set of anchor nodes and a much larger set of unidentified genotypes (such as an anonymized genotype database, e.g., UK Biobank). Without access to the latter set, the accuracy of identification of a random individual would be negligible, due to the small number of anchors. However, for a few of the unidentified genotypes, identification yields a candidate with a high confidence score (as we show, confidence scores can be used to accurately estimate probability of correct identification). The analyst adds to the set of anchor nodes and repeats the process.

## Results

As the number of anchors available to the analyst increases, so does the accuracy of identification, that is, the probability of correct identification of an anonymous genome belonging to a randomly sampled node. For example, the accuracy reaches 50% with as little as 1% of nodes being anchors for our default set of parameters (Table 1). While varying the population parameters and the error rates does have an impact on accuracy, the impact is small.

**Table 1:** Standard parameter values. Additional details in Supplement.

| Parameter | Explanation | Value |
| --- | --- | --- |
| Non-paternity rate | Fraction of nodes for which the analyst genealogy contains an incorrect father | 2.5% |
| Adoption rate | Fraction of nodes for which both parents are incorrect in the analyst genealogy | 0% |
| Missing parent rate | Fraction of parent-child links that are missing in the analyst genealogy | 5% |
| Number of islands | Number of islands in the island model of the population | 5 |
| Migration rate | Probability that an individual moves to a different island before mating | 20% |
| Monogamy rate | Fraction of individuals that produce offspring with only one mate | 50% |
| Number of generations | The number of generations in the god’s eye view of the simulation | 10 |
| Analyst horizon | The number of generations of genealogy that are available to the analyst | 6 |
| Identification horizon | The identification targets are sampled from the youngest few generations | 3 |

By outputting only high-confidence predictions, the analyst can achieve a substantial boost in precision (from 60% to 95% with the default set of parameters) with only a halving of recall (Figure S1). Alternately, by examining the top *k* predicted sibships, the analyst improves her chances that at least one of them contains the correct node: 75% for *k* = 10 and 84% for k=100 for the default set of parameters, compared to 60% for *k* = 1 (Figure S2).

Next, we seek to explain why identification works and how and when it fails. First, we see that for our standard parameters, a random individual has many IBD relatives among the set of anchors in the analyst view — mean: 5.05; median: 5 (Figure S5). The algorithm is able to successfully triangulate the individual based on multiple distant relatives: Figure S8 shows that the accuracy is over 50% even for targets whose closest anchor relative is at the 7th degree of relatedness. Younger generations have more anchor relatives than older generations, resulting in identification being more accurate for younger generations (72% for the youngest generation vs. 52% for the third-youngest generation with standard parameters). Another way to see that distant relatives are useful for identification is that accuracy drops substantially when the analyst horizon is curtailed: 31.6% for 5-generation horizon and 13.6% for a 4-generation horizon (compared to 60% for a 6-generation horizon).

When the algorithm fails, one common reason are the presence of error in the analyst’s view close to the target. For example, in sim 41% of the cases where the algorithm misidentifies, there is an error or missing link on the path from the target node to a grandparent. This compares with 22% in the cases the algorithm is correct). Another common failure case is when the algorithm identifies a close relative of the target instead: In 25% of error cases, the identified node is a cousin or closer relative of the target node (Figure S7). Finally, our assumption about the accuracy of IBD detection — specifically, the cutoff above which we assume that IBD segments are detected by the analyst — has little impact on accuracy (Figure S6).

Figure 3 shows the accuracy of snowball identification for our standard parameters (averaged over 10 runs of snowball identification). Only 300 nodes (0.1%) are anchors (note that our standard identification algorithm performs poorly with so few anchors). The analyst is given 30,000 unidentified dense genotype records (which corresponds to 10% of the population in the identification horizon). These genotype records are known to come from a set of 30,000 candidates nodes (for example, participants in a genetic study), but the mapping between genotype records and nodes is unknown to the analyst.

**Figure 3:**
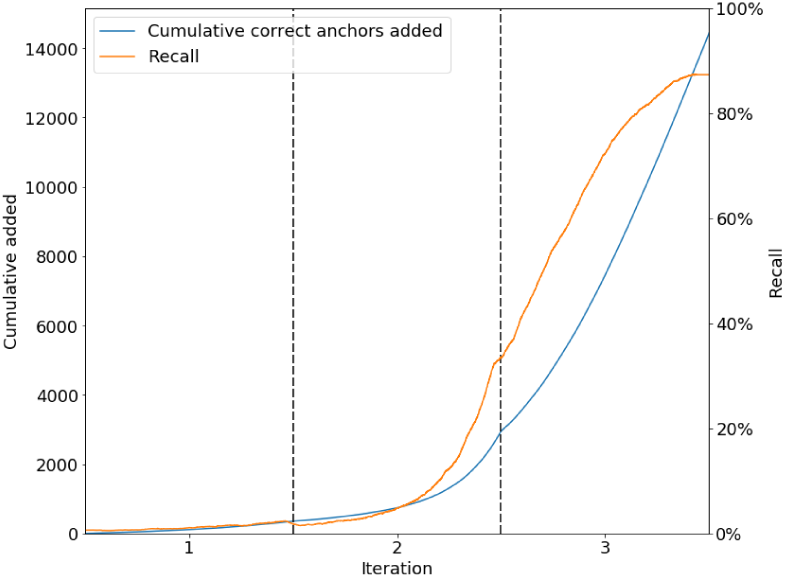
Cumulative correct and recall of snowball identification.

Since the analyst calibrates the identification algorithm to have high precision (95%), it initially has low recall (about 7%): that is, for most targets, the algorithm is not confident enough to make a guess. After iterating through the set of unidentified genotype records once (iteration count 30,000), the number of correctly identified records is about 360 and the number of incorrectly identified records is about 25. Since all identified genotype records/nodes are added to the anchors, at this point the analyst has about 680 anchors. By the end of the third iteration through the set of unidentified records, recall has improved to 90%; precision remains high throughout. All results in this paragraph are averages over 10 runs.

Snowball identification fails if the number of anchor nodes is too low (the higher the error, the higher the number of anchor nodes needed), or if the analyst calibrates the threshold too aggressively.

## Discussion

With proper oversight, identification by triangulation can be a powerful tool for law enforcement. One barrier to its use is the ability to extract whole-genome genotypes from degraded crime-scene samples, but techniques for doing so have improved substantially [10], leading to a quickly growing list of crimes solved through long-range familial searches (but not, to our knowledge, techniques resembling triangulation) [11].

Our techniques are directly applicable in a setting where the investigation is conducted with the cooperation of a consumer genetics company that maintains a large genealogical database, such as Ancestry Inc. Alternatively, investigators may be able to compile genealogical information from public and quasi-public sources such as vital records (birth, death, and marriage records) or the ancestral records maintained by the Church of the Latter Day Saints [12]. Finally, it may be possible to avoid the need for large-scale genealogy: investigators may upload the unknown genotype to a consumer genetics company such as GEDmatch, obtain a list of relatives, and only then construct genealogies of these relatives to look for intersections between the family trees. This approach would require modifications to the algorithm to account for the incompleteness of the genealogy, and we have not simulated its effectiveness. Nonetheless, it might be the most likely approach to be adopted in the near term, as it is a straightforward extension of the existing practice and avoids alerting the consumer genetics company to the provenance of the genotype data.

The usefulness of genealogical triangulation is not limited to law enforcement. DNA testing is already used by adoptees, children conceived by sperm donation, and victims of misattributed parentage to identify their biological parents; genealogical triangulation could substantially simplify these searches.

Genealogical triangulation can be seen as one example of a class of statistical inference techniques for reconciling two sources of genealogical information: recorded genealogy, which is typically precise but error-prone, and genetic genealogy, which is relatively accurate but stochastic. As such, techniques similar to ours may be useful to genetic genealogy services for detecting errors, merging family trees, and improving the accuracy of their predictions. They might also be useful to researchers for constructing pedigrees, especially of isolated populations [13].

On the other hand, the technique presents clear privacy risks to individuals and to society. Individuals adopting false identities, such as covert agents or people in witness protection, may be unmasked if adversaries are able to obtain a sample of their DNA. Further, research databases of genotypes are typically shared in a de-identified manner to protect participant privacy [9]; our findings provide a way to re-identify participants that is more accurate and broadly applicable than previous methods based on phenotypes or surname inference [14],[15]. For large de-identified databases, our snowball identification technique is applicable. More speculatively, if DNA amplification techniques continue to improve to the point that dense genotypes may be extracted from hair and other types of abandoned DNA, genealogical triangulation could be a route to mass surveillance in public spaces. As knowledge of these possibilities spreads, it might have a chilling effect on people’s willingness to contribute to research or participate in direct-to-consumer genetic testing.

So far, public policy on genetic privacy has been premised on secrecy and informed consent, with the goal of ensuring that no one can learn an individual’s genetic information without her knowledge and cooperation. Our work calls into question whether this approach will be sustainable in a world where a subset of individuals are willing to reveal their genotypes to others (sometimes even publicly), the cost of genotyping continues to plummet, and large-scale genealogical information is easy to obtain.

## Acknowledgement

This research was supported by the National Science Foundation under NSF Award IIS Award 1704444.

## Supplement

### Additional details of simulations

#### Populations and mating

We use an island model to synthesize a population. In this model, each individual is born in one of a fixed number of islands. Upon entering adulthood, each individual either stays on the island of their birth or moves to a different randomly selected island with a fixed transition probability. Individuals move between islands at most once. Individuals have 1–3 partners randomly selected from their island, with the exclusion of siblings (for the distribution of the number of partners, see the code) and an average of 2 offspring, so that there is no population growth over time. The population is divided into non-overlapping generations, and individuals in a mating pair belong to the same generation.

#### Genomes

We use an approach similar to Kessner et al. [8] to generate genomes. Each of the founders’ autosomes is assigned a unique identifier. We assume that no pair of founders shares IBD regions with each other. For descendants of the founders, contiguous regions of the genome are represented by the unique identifier of the founder that region of the genome is inherited from.

When generating a new genome, for each parent, we randomly select which of the homologous chromosomes the child will inherit. Recombination is simulated using the data provided by the HapMap project. The recombination rates provided by HapMap are averaged by sex; we used data from Kong et al. [16] to apply a scaling factor for males and females.

#### Twins

Approximately 1 in 333 births result in monozygotic twins, or a pair of individuals with identical DNA [17]. This is included in our simulations.

#### Errors in IBD detection

IBD detection is imperfect and produces false negatives — some IBD segments may not be detected, especially segments smaller than 5cM [18]. We assume that all segments longer than 5cM are detected and no segments shorter than 5cM are detected. Figure S6 shows that varying this threshold in the 0–10 centimorgan range has a negligible effect on the analyst’s accuracy.

##### Supplementary Algorithm 1: Identification with confidence score.

Given: a population genealogy, dense genotypes of anchors, and an anonymous genotype.

Let *node*_*i*_ refer to the *i*^*th*^ node ranked by LOGPROB (*node*) where LOGPROB is as defined in Algorithm 2.

Identification: Output the sibship containing *node*_*1*_.

Confidence score: Output LOGPROB(*node*_*1*_) – LOGPROB(*node*_*i*_) for the smallest *i* such that *node*_*1*_ and *node*_*i*_ are not in the same sibship.

---

**Supplementary Figure S1:**
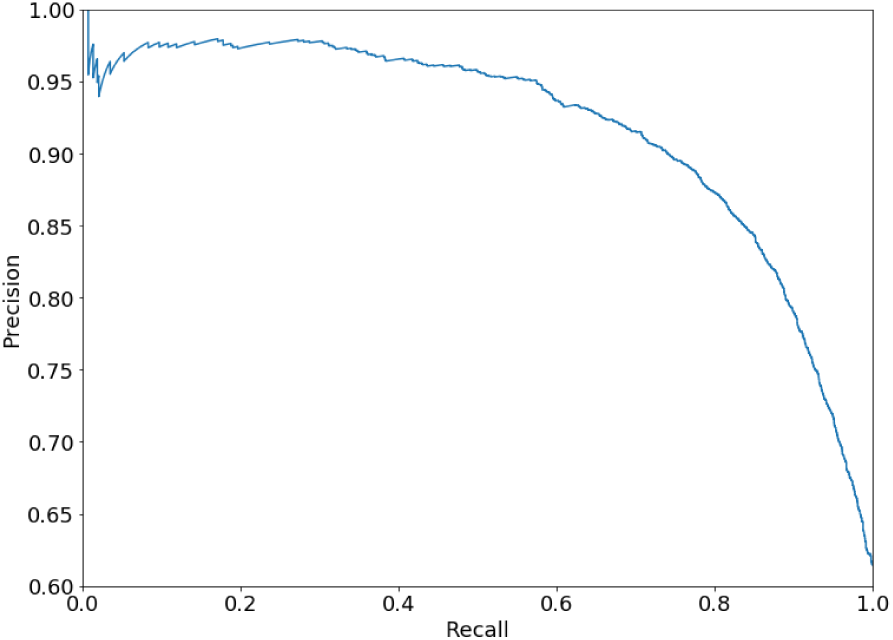
Trade-off between precision and recall using Supplementary Algorithm 1.

**Supplementary Figure S2:**
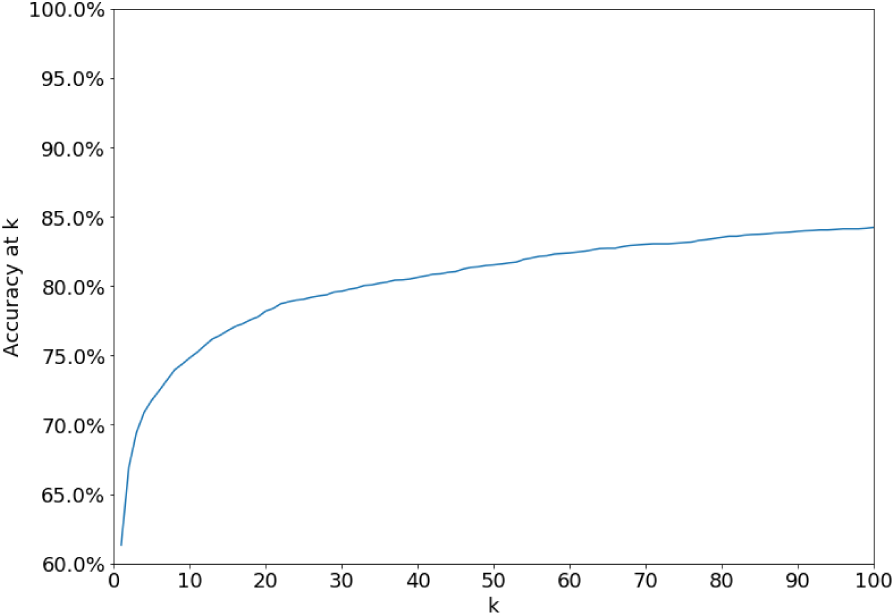
Accuracy as a function of *k* when identification is considered successful if the target individual is in one of the top *k* sibships.

**Supplementary Figure S3:**
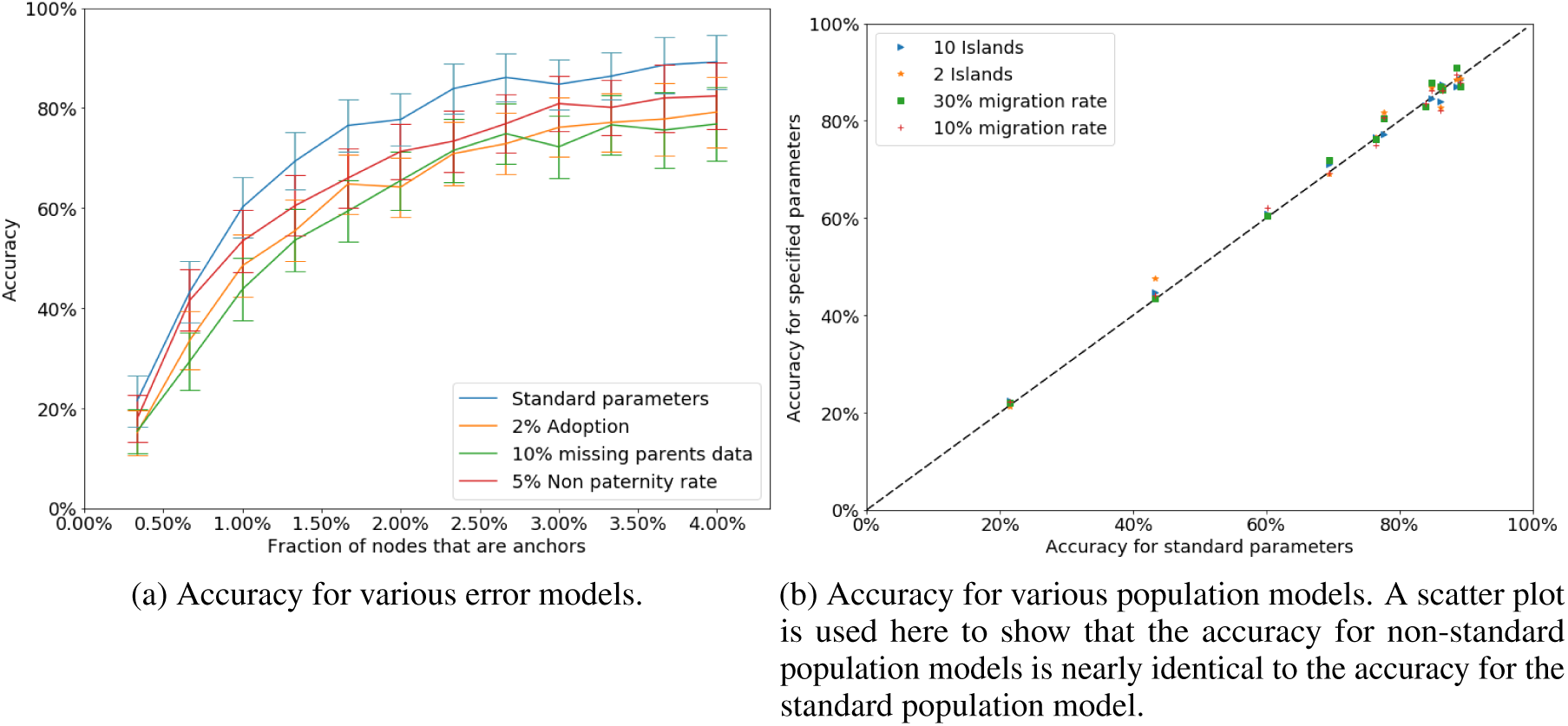
Accuracy for non-standard parameters.

**Supplementary Figure S4:**
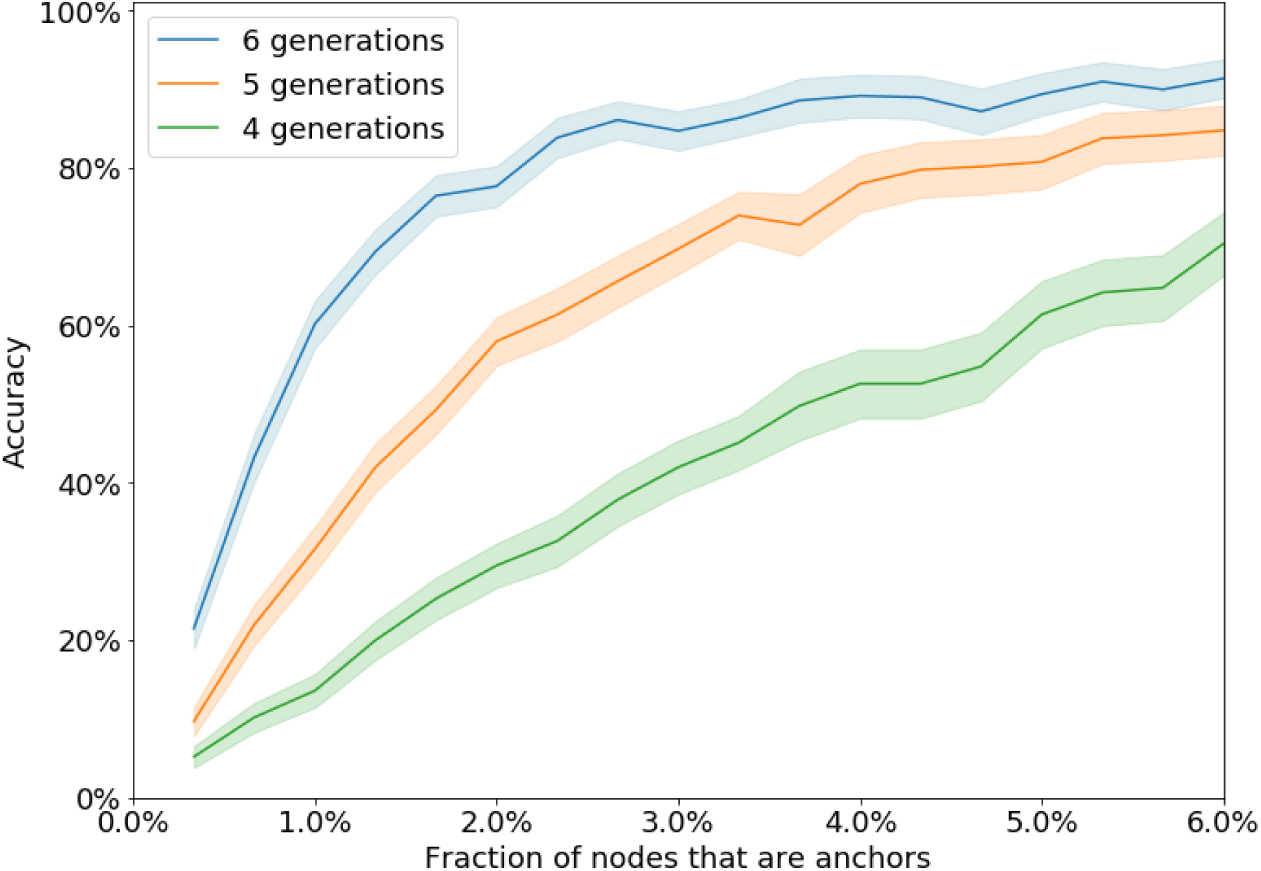
Accuracy for different values of the analyst horizon.

**Supplementary Figure S5:**
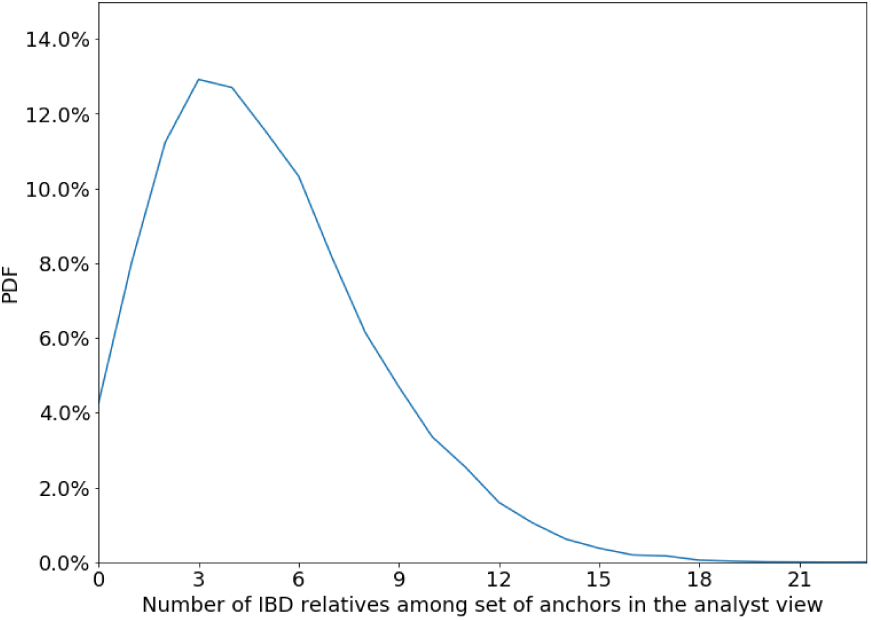
Distribution of number of anchors with IBD above 5cM that is predicted by analyst genealogy.

**Supplementary Figure S6:**
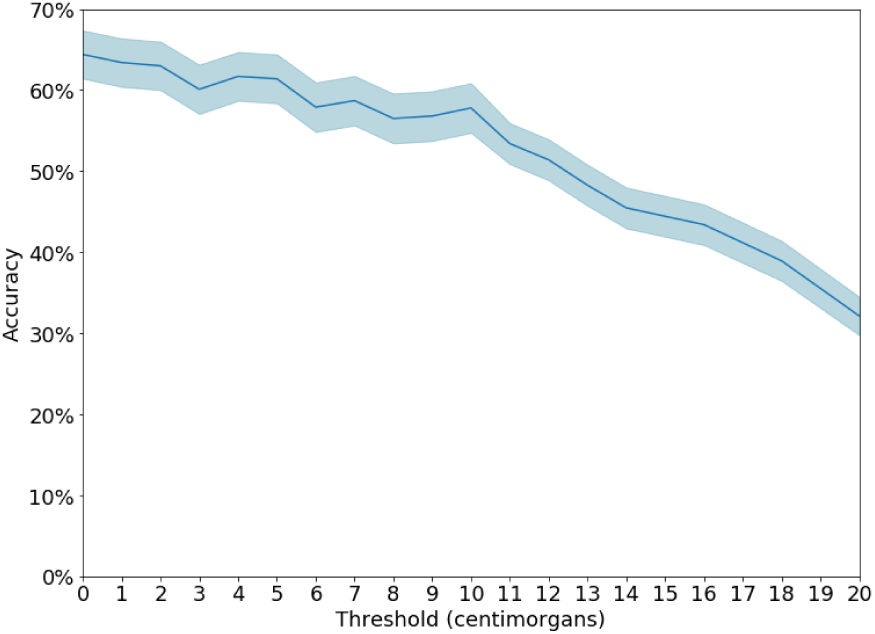
Accuracy as a function of IBD detection threshold.

**Supplementary Figure S7:**
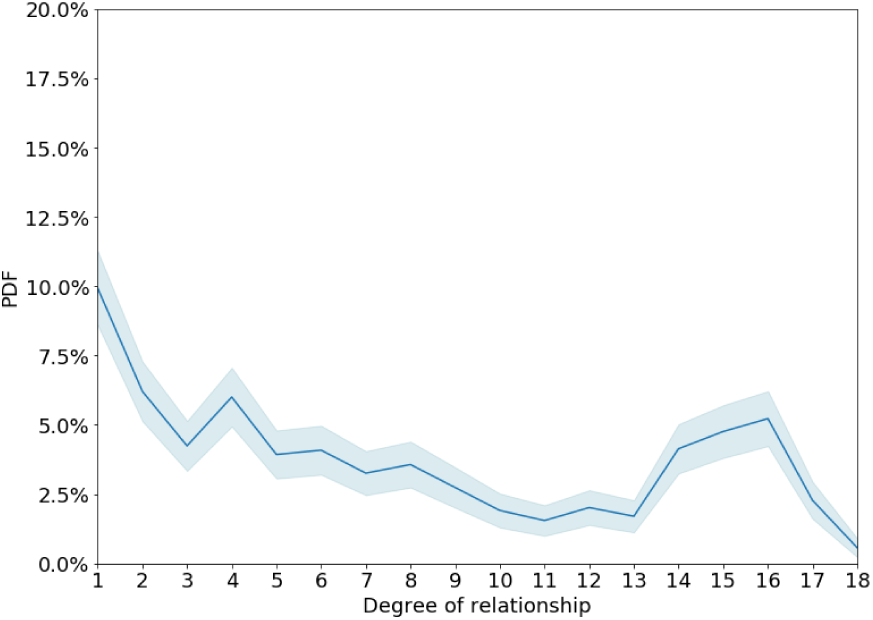
Degree of relationship between the target node and the incorrectly identified node in cases where identification is incorrect.

**Supplementary Figure S8:**
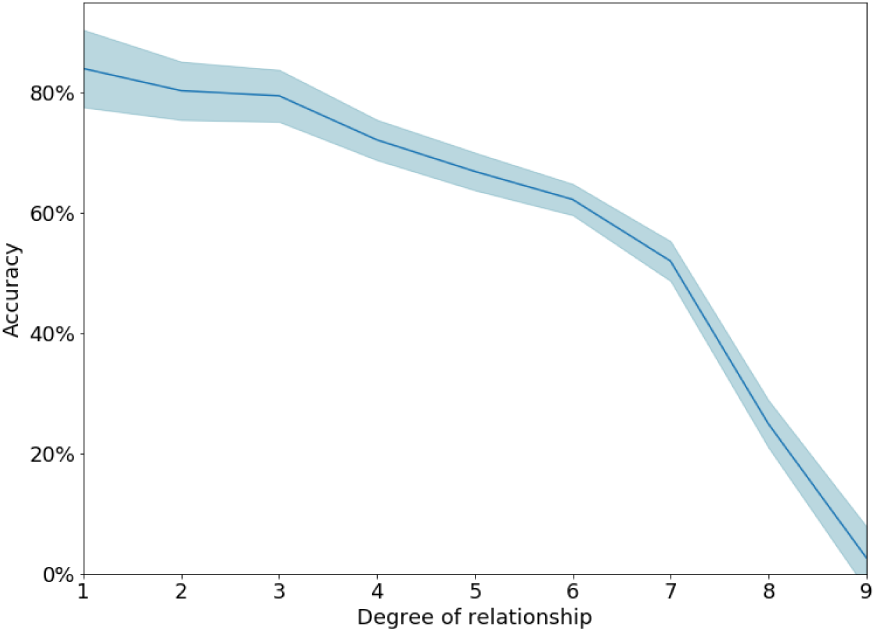
Accuracy as a function of the degree of relationship between the target node and the closest anchor node.

## Footnotes

1 The hurdle gamma distribution of IBD length is defined by two random variables. A Bernoulli random variable determines if the IBD length is zero or positive. If it is positive, then the value is drawn from a gamma distribution.

